# Predicting Lifespan-Extending Chemical Compounds with Machine Learning and Biologically Interpretable Features

**DOI:** 10.1101/2022.11.20.517230

**Authors:** Caio Ribeiro, Christopher K. Farmer, João Pedro de Magalhães, Alex A. Freitas

**Affiliations:** School of Computing, University of Kent, Canterbury, Kent, UK; Centre for Health Services Studies, University of Kent, Canterbury, Kent, UK; Genomics of Ageing and Rejuvenation Lab, Institute of Inflammation and Ageing, University of Birmingham, Birmingham, UK

**Keywords:** Machine learning, feature selection, lifespan-extension compounds, longevity drugs

## Abstract

Recently, there has been a growing interest in the development of pharmacological interventions targeting ageing, as well as on the use of machine learning for analysing ageing-related data. In this work we use machine learning methods to analyse data from DrugAge, a database of chemical compounds (including drugs) modulating lifespan in model organisms. To this end, we created four datasets for predicting whether or not a compound extends the lifespan of *C. elegans* (the most frequent model organism in DrugAge), using four different types of predictive biological features, based on compound-protein interactions, interactions between compounds and proteins encoded by ageing-related genes, and two types of terms annotated for proteins targeted by the compounds, namely Gene Ontology (GO) terms and physiology terms from the WormBase’s Phenotype Ontology. To analyse these datasets we used a combination of feature selection methods in a data pre-processing phase and the well-established random forest algorithm for learning predictive models from the selected features. The two best models were learned using GO terms and protein interactors as features, with predictive accuracies of about 82% and 80%, respectively. In addition, we interpreted the most important features in those two best models in light of the biology of ageing, and we also predicted the most promising novel compounds for extending lifespan from a list of previously unlabelled compounds.

## 1 Introduction

Old age is a major risk factor for a number of diseases, including many types of cancer, cardiovascular and neurodegenerative diseases [1] [2] [3]. Hence, there has been growing interest in developing interventions that target the biological process of ageing, in order to extend lifespan and healthspan [4] [5]. Non-pharmacological interventions like dietary restriction and genetic interventions have been quite successful for extending the lifespan of model organisms [6] [7] [8] [9]. However, genetic interventions tend to be unethical for humans, and arguably relatively few people would be willing to undergo dietary restriction for a long term. Hence, pharmacological interventions are currently the most promising type of anti-ageing intervention for extending human lifespan and healthspan, and this is current a very active research area in the biology of ageing [10] [11] [12].

A large number of compounds have been found by *in vivo* experiments to be able to prolong the lifespan of model organisms – in particular, the DrugAge database contains data on 1096 compounds that have been shown to extend the lifespan of model organisms [13]. Intuitively, the analysis of such data can lead to the discovery of novel lifespan-extending compounds, as well as potentially a further understanding of the underlying mechanisms of the biology of ageing [14] [15].

However, it is not feasible to manually analyse the relatively large volumes of data in DrugAge or other databases describing how each compound interacts with the biology of an organism. Hence, a promising research direction consists of analysing the data in such databases using machine learning algorithms that highlight patterns in data, particularly classification algorithms, which learn predictive models from data [16]. Therefore, recently there has been growing interest on applying classification algorithms to the data in DrugAge [17] [18] [19] [20], in order to learn models that predict which compounds are more likely to extend the lifespan of a given organism, which is also the overall goal of this work.

In this context, we have prepared four datasets with different types of features describing the properties of chemical compounds (including drugs) or proteins interacting with those compounds, and trained classification models using supervised machine learning (ML) methods to predict whether or not a compound significantly extends the lifespan of *C. elegans* worms, based on data in DrugAge and other databases.

The most widely used model organism for studying biological mechanisms of ageing is *C. elegans*. It has several characteristics that facilitate *in vivo* experiments and ageing research, such as being easy and inexpensive to maintain, having a short lifespan of about 3 weeks, being easy to observe and having fully sequenced genes that are in great part homologous to human genes [21]. In addition, of particular relevance to this work, they are the model organism with the largest amount of data in the DrugAge database (667 out of 1096 entries) [13].

The computational process of knowledge discovery goes from the collection and preparation of the data to the analysis of the patterns found by the ML algorithms. During the data preparation process, for datasets with a large number of features (also called variables), like our datasets, it is common to perform a pre-processing task of feature selection (FS), which involves analysing the relationships in the data to select the most relevant features (independent variables) for the task at hand. We focused on the task of FS in this study, more specifically on filter methods, which rank features based on their relationship to the classification task, i.e., how much they influence the value of the class (target) variable. Hence, we applied filter methods in a pre-processing phase and used the selected features to train a classifier using the well-known random forest classification algorithm [22]. The most accurate predictive models were then further analysed to identify the features most relevant for our classification task and to identify novel compounds which have a high probability of extending the lifespan of *C. elegans*, as estimated by those best models.

The datasets used in our experiments were generated following a methodology which is broadly similar to the one used in the study by Barardo et al. [17]. In their study, the authors used Random Forest (RF) classifiers for the same prediction problem, also using *C. elegans* DrugAge data to obtain the instances (ageing-related compounds). That study, similarly to other related works ([19], [18]), uses a combination of GO Terms and chemical descriptors of the compounds, applying ML techniques to the same problem of discovering compounds related to *C. elegans*’ longevity.

The novel contributions of our study compared to the related work in [17] [18] [19] [20] are as follows. First, we created datasets using four different types of predictive features, based on Gene Ontology (GO) terms, drug (compound)-protein interactions, interactions between compounds and proteins encoded specifically by ageing-related genes, and physiology terms from a Phenotype Ontology for *C. elegans*. GO terms have been used as predictive features in [17] [18], but the other three types of features proposed here are new types of features for predicting a compound’s effect on the lifespan of an organism using machine learning, to the best of our knowledge. Note that we do not use chemical descriptors as features, an approach used in [17] [18] [19] [20], which generated models with good predictive accuracy. We do not use chemical features because they represent very specific chemical information which is not very meaningful for biogerontologists. For instance, the three most important molecular descriptors in the best model learned in [19] were ‘number of nitrogen atoms’, ‘total positive van der Waals surface area of atoms with a partial charge in the range of 0.10 to 0.15’, and ‘hydrophobic volume’; which do not shed light on the kind of biological process associated with a drug. Hence, instead of such chemical descriptors, we use only biologically interpretable features, representing potentially relevant information for biogerontologists.

The second contribution of this study is the proposal and evaluation of an approach that automatically selects the best filter method for feature selection from a set of 5 candidate filter methods, using the training data. Finally, as additional contributions, we also perform a biological analysis of the most important predictive features in the best (most accurate) classification models learned from our datasets, and identify the most promising new compounds in our dataset for extending *C. Elegans* longevity – i.e., compounds that are annotated in the dataset as not extending *C. elegans*’ lifespan but which are predicted by our best models, with a very high probability, to be capable of extending the lifespan of *C. elegans*.

## 2 Results and Discussion

As mentioned, our experiments were performed on four datasets prepared for this study, all based on data from the DrugAge database and other sources (see Section 4.1). In these datasets, each instance (record) represents a compound (drug), which consists of a set of predictive features (variables) and a class label to be predicted. The class labels indicate whether or not a compound was found to significantly extend *C. elegans*’ lifespan, represented as a positive or negative class label, respectively. In essence, a compound is assigned a positive class if there is an entry in the DrugAge database [13] showing that the compound extended the average lifespan of C. elegans by at least 5% and the extension was statistically significant, whilst the list of negative-class compounds was obtained mainly from related work [17], consisting of compounds which do not satisfy the above criterion for the positive class (see Section 4.1 for a more precise definition of the negative class).

The four created datasets have approximately the same instances (compounds) and class labels, but each dataset has a different set of binary predictive features, as graphically summarised in Figure 1.

**Figure 1.**
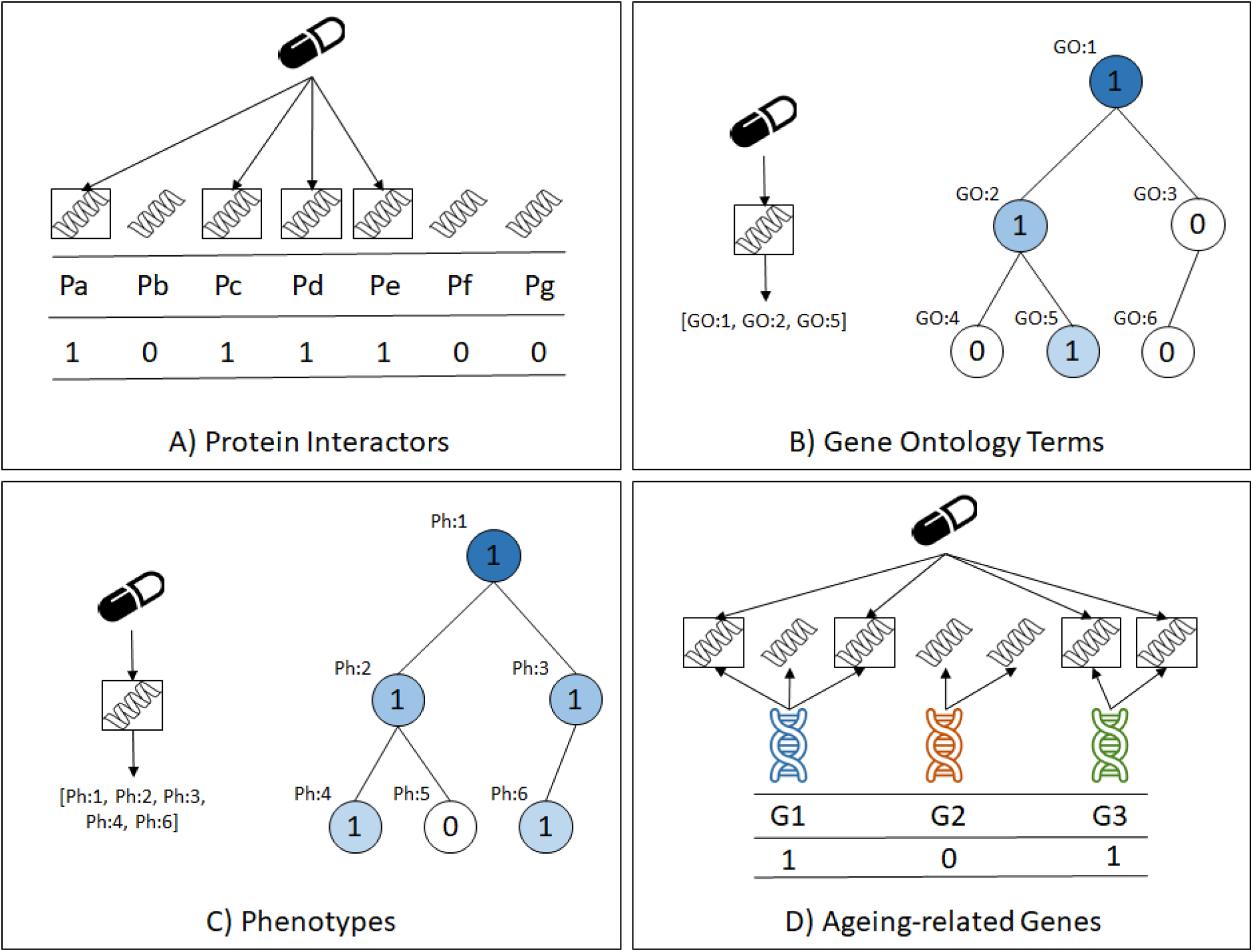
The four types of predictive features in the datasets created for this study

More precisely, the four types of features (datasets) are: (a) *Protein interactors*: in this dataset each feature represents a protein and takes the value 1 or 0 to indicate whether or not that protein interacts with the compound associated with the current instance; (b) *Gene Ontology* (*GO*) *term annotations*: in this dataset each feature represents a GO term [23] and takes the value 1 or 0 to indicate whether or not the compound associated with the current instance interacts with at least one protein that is annotated with that GO term; (c) *Physiology phenotype annotations*: in this dataset each feature represents a physiology term from the WormBase Phenotype Ontology [24], and takes the value 1 or 0 to indicate whether or not the compound associated with the current instance interacts with at least one protein that is annotated with that physiology term from the Phenotype Ontology; and (d) *Ageing-related genes*: in this dataset each feature represents a gene in the GenAge [25] or the GenDR [26] database, and it takes the value 1 or 0 to indicate whether or not the compound associated with the current instance interacts with at least one protein encoded by a gene in GenAge or GenDR.

Note that the first feature type, protein interactors, represents a ‘direct’ property of a compound (drug), whilst the latter three feature types represent ‘indirect’ properties of a compound, in the sense that they are associated to the protein interactors and, by extension, to the compounds. To the best of our knowledge, of these four feature types, only GO Terms have been used for predicting lifespan-extension compounds with machine learning algorithms [17] [18], and the other three feature types are novel contributions in this context.

Note that, in Figures 1(B) and 1(C), the GO term annotations and the physiology phenotype annotations datasets have features that are hierarchically related via a generalization-specialisation relationship. That is, if an instance (compound) takes the value 1 for a GO term or physiology phenotype feature, that instance will also take the value 1 for ancestors of that GO term or phenotype feature in the corresponding hierarchy, where the ancestors represent more general properties than their corresponding descendant features in the hierarchy.

The datasets have large numbers of features in general: 5788 features in the protein interactions dataset, 7706 in the GO terms dataset, 1272 in the physiology phenotype dataset, and 365 in the ageing-related genes dataset. Therefore, we applied feature selection methods to each dataset in a data pre-processing phase, before using a classification algorithm to learn a predictive model from the data. We performed experiments comparing six different feature selection (FS) methods. Five of these are based on well-established FS methods that follow the filter approach [27] [28], where a FS method measures the degree of association between each feature and the class variable and then selects the top-ranked features based on those measures – see Section 4.2 for details. The experiments also included ensemble versions of those filter methods, called filter ensembles, which combine the outputs of many runs of a filter method in a way that mitigates the problem of class imbalance in our datasets in order to improve robustness and predictive accuracy, as explained in Section 4.3. The sixth type of FS method is a novel FS approach proposed in this paper (Section 4.5), called Auto-Filter, that automatically performs a data-driven selection of the best filter or filter ensemble method for the input data, from a set of pre-defined candidate filter methods (Section 4.2) or filter ensemble methods (Section 4.3). The Supplementary Material for this paper has the detailed results of this comparison of FS methods, which resulted in the proposed Auto-Filter being selected as the best filter approach.

### 2.1 Predictive Accuracy Results

In this Section we report on the predictive accuracy results obtained with the following experimental setup. The classification models were trained using a Random Forest (RF) algorithm, which is among the top-performing classification algorithms in general [29] [30] and is very popular in bioinformatics, and has also been used in previous studies for predicting lifespan-extension compounds [17] [18] [19]. A random forest algorithm also has the advantage of facilitating an indirect analysis of feature importance, which can be useful for detecting highly predictive features for a given classification problem, as shown later. More specifically, we used the ‘Balanced Random Forest’ method [31] to cope with the class-imbalance issue (see Section 4.6 for details).

As the RF algorithm has an embedded feature selection process, it is considered to be robust against datasets with a large number of features (like our datasets), so it is possible that performing FS in a data pre-processing step would not have a positive impact on the predictive accuracy of the resulting RF classifier. Therefore, we compared the results of a RF classifier trained using our proposed filter approach, Auto-Filter, against a Baseline RF classifier using all the original features, i.e. not performing any FS prior to training the classifier.

The predictive accuracy of the learned Random Forest classifiers was measured by the popular Area Under the ROC curve (AUC) measure, using a standard 10-fold cross-validation procedure [32]. The AUC measure takes values in the range [0..1], where 0.5 is the expected score for randomly guessing the class labels and 1 would be the score of a perfect classifier. We report the median AUC results over the 10 test folds of the 10-fold cross-validation, since the median is more robust to outliers than the mean. The median AUC values of the Random Forest classifiers trained with the features selected by the different filter ensemble methods are reported in Table 1. The highest AUC value for each dataset is highlighted in boldface font in this table.

**Table 1.** Median AUC results for RF trained with the features selected by the Auto-Filter approach and the baseline RF trained with all features (i.e., no feature selection in a pre-processing step)

| Dataset | Baseline | Auto-Filter |
| --- | --- | --- |
| Protein Interactors | 0.717 | **0.801** |
| GO terms | 0.767 | **0.818** |
| Phenotypes | **0.741** | 0.728 |
| GenAge/GenDR | 0.683 | **0.725** |

For this comparison, we performed a statistical analysis to investigate whether the use of the Auto-Filter approach significantly outperforms the baseline of no feature selection in a pre-processing step. We applied a two-tailed Wilcoxon signed-rank test [33] (a non-parametric test that does not make assumptions about the data distribution) comparing the AUC results of each fold of the 10-fold cross-validation process pairwise, meaning the same training and test folds were used for each pair of classifiers we compared. Note that this means that the sample size for the Wilcoxon signed-rank test is 10, corresponding to the results on the 10 test sets of the 10-fold cross-validation procedure.

None of the 4 comparisons wielded significant results (p-values *0.275, 0.625, 0.921* and *0.921* for Protein Interactors, GO Terms, Physiology Phenotypes and GenAge/GenDR datasets respectfully), meaning we could not reject the null hypothesis that the methods’ performances are statistically equivalent.

However, considering that the difference between the median AUC results was relatively high in some cases in Table 1 (e.g., 8.4% for the Protein Interactors dataset), it is possible that the small sample size (10) used by the Wilcoxon signed-rank test had too much influence on the results of the statistical test, since the p-values computed by statistical significance tests are quite sensitive to the sample size, and lack of statistical significance does not mean lack of biological relevance [34] [35] [36] [37] [38]. Thus, we also calculated a measure of the ‘effect size’ for the differences of AUC values between the two methods in Table 1, which is a measure much less sensitive to sample size and is thus more suitable for identifying differences that are relevant in practice [39] [34], particularly when using a small sample size.

More precisely, we calculated the popular Cohen’s *d* measure of effect size [34] for each dataset, as an approach for investigating the difference between the AUC values of the Baseline RF and the RF with Auto-Filter, for each dataset (feature type). The effect sizes are usually classified into small (*0*.*2* ≤ *d < 0*.*5*), medium (*0*.*5* ≤ *d < 0*.*8*) and large (*d ≥ 0*.*8*), reflecting how apparent the difference between the groups is (with *d < 0*.*2* indicating an irrelevant or negligible effect). For our comparisons of the results in Table 1, we found *d* = 0.617 for the Protein Interactors dataset (a medium effect) and *d* = 0.291 for the GO Terms dataset (a small effect), which indicate a relevant difference in AUC values for both datasets. Regarding the other two datasets, we got effect sizes of *d* = 0.051 for Phenotypes and *d* = 0.027 for GenAge/GenDR, both indicating that the differences in the AUC values of the two approaches are negligible.

Based on these results, we can conclude that Auto-Filter was superior to the baseline approach, as it had the best median AUC for 3 out of 4 datasets, and the only two relevant differences in AUC values according to the effect size analysis were in favour of the Auto-Filter approach.

Recall that even though the predictive features in the 4 datasets are different, they share approximately the same instances (compounds) and the same prediction problem (i.e., the same class label). Therefore, it is interesting to discuss the effect of the type of feature on the predictive accuracy of the classifiers.

Across all RF classifiers trained in our experiments (including those trained in our preliminary experiments comparing different FS methods, see Tables S1-S4 in the Supplementary File), the ones with the highest median AUC values were the RF classifiers trained with the features selected by the Auto-Filter approach for the Protein Interactors (AUC = 0.801) and GO Terms (AUC = 0.818) datasets. We selected these two classifiers for an analysis of their most important predictive features (Section 2.2). In addition, we selected the RF classifier trained with the features selected by Auto-Filter for GO terms (the most accurate classifier overall) for identifying the most promising novel compounds for *C. elegans*’ lifespan extension (Section 2.3).

### 2.2 Analysis of Feature Importance in the Best Predictive Models

Supervised machine learning models, in addition to being tools for predicting target variables, reflect patterns in the data used to train them. Interpreting a classification model, by identifying how its predictions are made, can help the user both check the internal consistency and biological validity of the decisions made by the algorithm and find interesting patterns that may spark new research directions.

Decision tree-based classifiers are particularly interpretable, as decision trees can be visualized and interpreted without the need of any other tools. In the case of Random Forest models, however, directly interpreting each random tree in the forest is not feasible, due to the large number of trees. As an alternative, we can calculate a feature importance measure, which allows the user to see which features are considered most important for classification across all trees in the forest. In this Section we analyse the top features of our two most successful predictive models, the Random Forests models trained with the features selected by Auto-Filter on two datasets: the Protein Interactors (median AUC = 0.801) and the GO Terms (median AUC = 0.818) datasets. For the latter model, we selected features from the Molecular Function and Biological Process categories, excluding Cellular Component features because we found that the selected top features in this category were redundant with respect to one of the top Molecular Function terms.

For this feature importance analysis, we trained new models using the entire datasets (no training and test set division), to ensure the model-interpretability analysis would consider all data available. The Balanced Random Forest method was used again to deal with class imbalance, in order to avoid a bias in favour of the majority class in the predictions.

The metric of feature importance used in this analysis was the Gini Importance Measure (GIM). The GIM is the average reduction in Gini Index over all nodes in the decision tree which use that feature for branching, over all decision trees in the forest [40], [41]. The main drawback of this metric in general is a bias in favour of features with many possible values. However, this drawback does not occur in our case as all our predictive features are binary. Tables 2 and 3 show the top 10 features based on GIM for the Protein Interactors and GO Terms (respectively) in the classification models learned by Random Forests with the Auto-Filter approach. Hence, the features in Tables 2 and 3 are essentially the features that are most relevant for discriminating between positive-class (lifespan-extending) and negative-class (non-lifespan-extending) compounds.

**Table 2.** The most important features in the RF classifiers trained with Protein Interactors as features and feature selection performed using the Auto-Filter approach.

| **Protein**<br>**(STITCH id)** | **Name** | **Positive**<br>**Class**<br>**Proportion** |
| --- | --- | --- |
| 6239.C44B7.7 | Gamma-glutamylcyclotransferase | 90%<br>(18/20) |
| 6239.ZK892.4 | CaiB/baiF CoA-transferase family protein<br>ZK892.4 | 94.1%<br>(16/17) |
| 6239.T06D8.8.1 | Proteasome Regulatory Particle, Non-ATPase-<br>like | 88.9%<br>(16/18) |
| 6239.Y47D9A.1a | Ntp_transferase domain-containing protein | 93.3%<br>(14/15) |
| 6239.T19E7.2a.2 | Nuclear factor erythroid 2-related factor 2;<br>Protein skinhead-1 | 93.3%<br>(14/15) |
| 6239.F35G12.12.1 | Not available (uncharacterised protein) | 87.5%<br>(14/16) |
| 6239.F49C12.8.1 | 26S proteasome non-ATPase regulatory subunit<br>6 | 87.5%<br>(14/16) |
| 6239.T03D8.3 | 7B2 (Mammalian 7BT prohormone convertase<br>chaperone) homolog; | 84.6%<br>(11/13) |
| 6239.C53H9.1.2 | 60S ribosomal protein L27 | 84.6%<br>(11/13) |
| 6239.T28D9.2a | Probable splicing factor, arginine/serine-rich 5 | 83.3%<br>(10/12) |

**Table 3.**
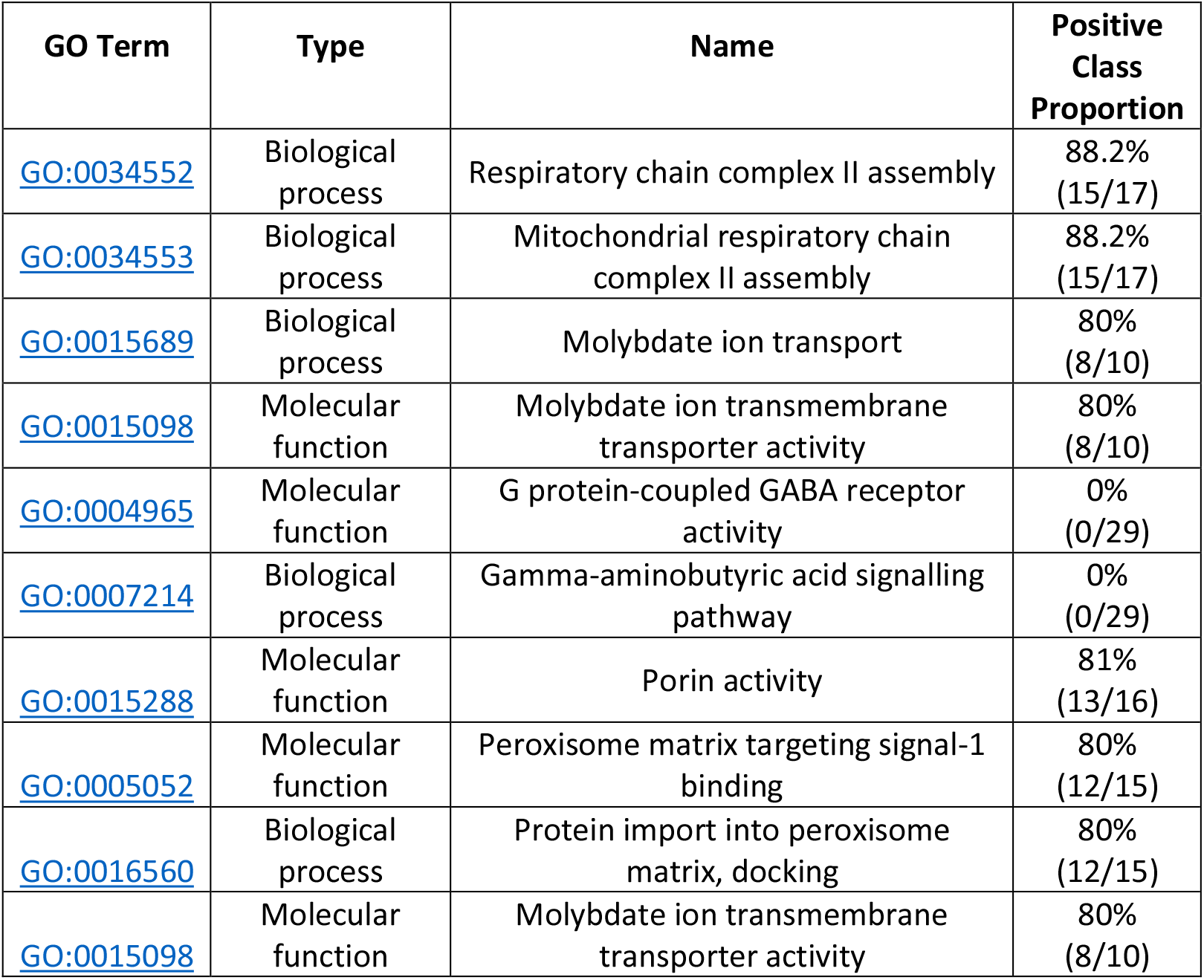
The most important features in the RF classifiers trained with GO Terms as features and feature selection performed using the Auto-Filter approach.

In these tables, the last column shows the proportion of positive-class instances (i.e., lifespan-extending compounds) among the instances which take the value ‘1’ for the feature – i.e., among all compounds annotated with the corresponding protein interactor or GO term. For example, for the first row in Table 3, there are 17 compounds annotated with the GO term ‘Respiratory chain complex II assembly’, out of which 15 (88.2%) are positive-class compounds. Note that there are two complementary ways for a feature to be one of the most relevant features (and so be included in Table 2 or 3): the feature can be a strong predictor of the positive class (i.e., the positive-class proportion is very high) or a strong predictor of the negative class (i.e., the positive-class proportion is very low). Most of these top features in Tables 2 and 3 are associated with the positive-class, but two features in Table 3 are instead associated with the negative-class – i.e., compounds taking the value ‘1’ for such features are in general negative-class compounds (their positive-class proportion is 0).

The top protein interaction (Table 2) and GO (Table 3) features associated with longevity drugs reflect processes commonly associated with aging and those more often targeted in longevity pharmacology. Gamma-glutamylcyclotransferase is the top feature (Table 2), an enzyme involved in glutathione metabolism [42] that, as an antioxidant, has often been targeted pharmacologically by longevity biotechs [12] [15]. Likewise, NRF2 (nuclear factor erythroid 2-related factor 2) and skn-1 (protein skinhead-1) are present in the GenAge database of age-related genes and have been previously associated with aging in model organisms, including worms [43]. The presence of several proteasome proteins as top interactors (Table 2) is also noteworthy and in line with several lines of work indicating that the proteasome is important for longevity and a promising target for longevity [15]. The mitochondrial respiratory chain complex is the top GO term (Table 3), reflecting that this complex has been associated with aging and is often targeted in longevity pharmacology [15] and showcasing the biological relevance of our results. Other GO terms, like porin activity, are also likely related to mitochondria. Therefore, and as expected, our results largely include processes and proteins previously associated with aging. In addition, we found terms (e.g., “G protein-coupled GABA receptor activity” and “Gamma-aminobutyric acid signalling pathway”) associated with the negative class; that is, terms predictive that a drug will not extend lifespan.

### 2.3 Identifying the Most Promising Novel Compounds for Lifespan Extension

In this Section we identify the 10 compounds, from a list of 5920 unlabelled compounds from DrugBank, with the highest probability of being classified as members of the life-expansion class (positive class) by our best classifier, the classifier trained with the GO Term features (with AUC = 0.818). For this task, we use only this classifier because it achieved the highest predictive accuracy in our experiments. The dataset of unlabelled compounds was created using the DrugBank database (version 5.0, downloaded in June 2022) [44], an online database of drugs and drug targets. The compounds selected in this analysis represent potentially novel compounds for extending *C. elegans*’ lifespan, although whether or not they really have this effect needs to be validated by proper biological experiments, of course, which is left for future research. The 10 top most promising compounds for our best classifier are listed in Table 4.

**Table 4.**
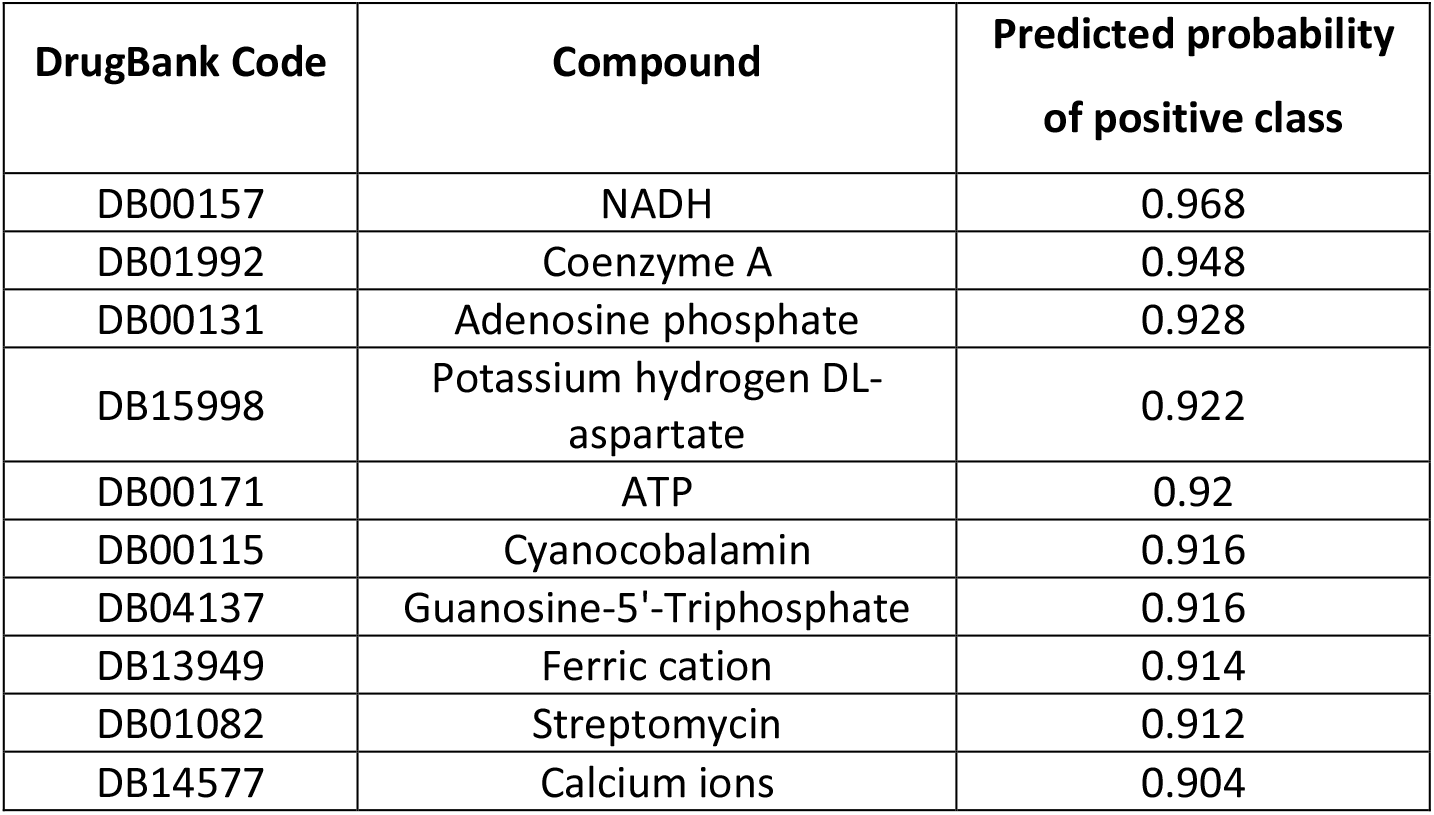
The 10 compounds from DrugBank with the highest probability of being novel compounds for extending C. elegans’ lifespan, from the best RF model trained with GO Term features

As a secondary criterion for selecting promising novel compounds, we measured the similarity of each of the top compounds in Table 4 (from DrugBank) to the compounds in our original dataset. This allowed us to determine how often their ‘neighbours’ (i.e., the most similar compounds in our dataset regarding the GO Term annotations of their protein interactors) are labelled as lifespan-extension compounds (positive-class). We used the Jaccard coefficient [45] to calculate the similarity between compounds – a measure of similarity between binary sets that only considers positive matches (ignoring matches of ‘0’ values), widely used in bioinformatics studies.

This criterion was chosen because, intuitively, the DrugBank compounds with many positive-class neighbours in our dataset, in addition to having a high probability of positive-class prediction by our best classification model, are the best choices for possible novel compounds for longevity research. Thus, we set a cut-out point of at least 80% positive-class neighbours, from the 20 most similar compounds in our original dataset, as our second selection criterion in this analysis. Based on this we selected four compounds to focus on: NADH (16/20 neighbours belonging to the positive class), Potassium hydrogen DL-aspartate (17/20), Ferric cation (17/20) and Streptomycin (17/20). The other 6 compounds in Table 4 have between 65% and 75% positive-class neighbours and, although still relevant as possible novel compounds, will not be discussed in detail.

The top predicted new longevity compound is NADH, the reduced form of NAD+, that is involved in metabolism and redox reactions. There has been significant interest into NAD+ and aging, including into NAD+ enhancers as a potential therapy [15] [46].

Interestingly, potassium hydrogen DL-aspartate has not, to our knowledge, been studied in the context of longevity; but has been shown in cells to inhibit damage and apoptosis from oxidative stress [47], and thus may be interesting to study in the context of longevity. Sun et al. [47] suggests that L-aspartic acid potassium salt protects from apoptosis and damage. This compound is chiral, meaning that there are two mirror images (isomers) of that same compound available. This means that in an environment where there are chiral targets (e.g., proteins) or other chiral molecules, this can affect properties that the molecule may have. Our best classifier suggests that Potassium hydrogen DR-aspartate (a mixture of both isomers) may have a role to play in longevity supported by this data on L-aspartic acid potassium salt (a single isomer of the same compound).

Ferric cation also, to our knowledge, has not been studied in the context of aging, although is it is interesting to note that iron metabolism has been associated with aging [48].

Also noteworthy, streptomycin is an antibiotic often used in *C. elegans* culture. If streptomycin were to extend lifespan in worms then it could be a potential source of bias in longevity studies, hence further studies are warranted.

## 3 Conclusions

We created four datasets for predicting whether or not a compound extends the lifespan of *C. elegans*, using data from the DrugAge database to assign a positive or negative class label to each compound, depending on whether or not the compound is recorded in DrugAge as significantly extending *C. elegans*’ lifespan by at least 5%. The datasets use four different types of predictive features, based on compound-protein interactions, interactions between compounds and proteins encoded specifically by ageing-related genes, and two types of terms annotated for proteins targeted by the compounds, namely Gene Ontology (GO) terms and physiology terms from the WormBase’s Phenotype Ontology. To analyse these datasets we used a combination of feature selection methods in a data pre-processing phase and the well-established random forest algorithm for learning a predictive model from the selected features.

The two best models were learned using GO terms and protein interactors as features, with a predictive accuracy of about 82% and 80%, respectively. In addition, we used a feature importance measure to identity the most relevant features in the best random forest models. Among those top-ranked features, there are several GO terms and protein interactors that are known to be associated with the ageing process, particularly involving the mitochondria and the proteasome, which are already promising targets for longevity drugs.

Furthermore, we identified the most promising novel compounds for extending *C. elegans* based on the predictions of the best learned random forest model – i.e., compounds from the DrugBank database (not included in the data used to train the classifiers) that were predicted with a very high probability to be positive class (extending lifespan) compounds. The most promising novel compounds (which have not been investigated in the context of ageing yet, to the best of our knowledge) include Potassium hydrogen DR-aspartate and streptomycin. Potassium hydrogen DR-aspartate is a mixture of two isomers, and the hypothesis of its potential pro-longevity effect in *C. elegans* is supported by the data for a single isomer of that compound (L-aspartic acid potassium salt). Streptomycin is an antibiotic often used in *C. elegans* culture, and so, if further research confirms that this compound really extends the lifespan of *C. elegans in vivo*, this would show an important source of currently undetected bias in longevity experiments with *C. elegans*.

Future research will involve lab experiments with *C. elegans* in order to try to confirm these computational predictions.

## 4 Methods

### 4.1 Dataset Preparation

We created four datasets, all with approximately the same instances (representing chemical compounds, or drugs) and the same definition of positive and negative class labels, but different types of predictive features (variables). The positive-class instances consist of drugs or compounds whose administration led to a statistically significant average increase of at least 5% of *C. Elegans*’ lifespan, as recorded in the DrugAge database (Build 4) [13]. The DrugAge database collects information of potentially life-extending compounds, based on publications reporting wet-lab experimental results (Website: genomics.senescence.info/drugs).

The list of negative-class instances (i.e., compounds found to have no significant positive impact on *C. elegans*’ lifespan) was taken mainly from the supplementary material provided in a previous study [17] that created a similar dataset, with two extensions, as follows. First, some compounds were included in DrugAge for having lifespan-increasing effects on other organisms, but their impact on *C. Elegans*’ lifespan was negative, so these compounds were used as negative-class instances. Second, some of the negative-class instances from the list in [17] were updated as positive-class based on more recent information in DrugAge, as the previous list was based on information from 5 years ago.

As mentioned earlier, we used four types of predictive features (one for each of the 4 datasets), namely features based on Gene Ontology (GO) terms, drug (compound)-protein interactions, interactions between compounds and proteins encoded specifically by ageing-related genes, and physiology terms from a Phenotype Ontology for *C. elegans*. Thus, all predictive features in our datasets are related to the proteins that interact with each compound, namely: the protein interactors themselves, the GO Term annotations and the Physiology Phenotypes associated with those interacting proteins, and whether the protein interactors of a compound are coded by an ageing-related gene. Notably, two of the four most related works [17] [19] [18] [20] (applying machine learning to DrugAge data) also use GO Term features, namely [17] and [18], but none of those four works used protein interactors, phenotypes or ageing-related genes as predictive features.

Our source of protein-compound interaction was the STITCH database (version 5.0, downloaded in 11-2021; Website: http://stitch.embl.de/) [49], a database of interactions between chemicals and proteins. We discarded all compounds that either were not found on STITCH or did not have any information of protein interactions stored there. In particular, we removed from our initial list of compounds (instances) all the entries for plant extracts that are not used commercially as drugs, since there is no entry in STITCH for such extracts. This filtering process caused our sample size to be reduced by about 25% compared to the original dataset in [17], arguably making the prediction problem more difficult. However, this was necessary for the types of predictive feature used in our datasets, as they are all based on protein interactors, and to compensate for the dataset reduction we got the benefit of creating datasets where all features are biologically interpretable, as mentioned earlier. After this instance (compound) filtering process, the datasets used in our experiments share approximately the same set of 1120 instances, 253 (22.6%) of which refer to lifespan-extending (positive-class) compounds in *C. Elegans*, with the remaining 867 (77.4%) being negative-class compounds. With this set of instances, we created four datasets based on different types of biological information, as described in the next four subsections.

Note that, after the initial creation of each dataset, we applied a simple frequency-threshold filter to remove features with fewer than 10 instances with a ‘1’ value in the training set (all features in our datasets are binary, with ‘1’ indicating the presence of the feature). This frequency-threshold filter was applied to reduce the risk of overfitting, and it was applied at the start of the data pre-processing phase, i.e. before applying the feature selection filters. In addition, if the application of this frequency-threshold filter results in an instance having all its features taking the value ‘0’ (rather than ‘1’), that instance is also removed from the dataset. As a result, there is a small variation in the number of instances in each of the four datasets, as follows: the Protein Interactors and GO terms datasets have 1120 instances, the Phenotypes dataset has 1103 instances, and the GenAge dataset has 1034 instances.

#### 4.1.1 Protein Interactors Dataset

For this dataset, we created binary features that indicate whether or not a protein interacts with the current instance (compound). The created dataset initially had 9880 unique protein interactors obtained from the STITCH database (version 5.0), which were reduced to 5788 predictive features, after applying the aforementioned simple frequency-threshold filter to avoid overfitting. The number of interactors associated with a given compound varies greatly, reaching over 1000 interactors for some well-known compounds.

#### 4.1.2 Gene Ontology Dataset

Expanding on the information from the previous dataset, we used the Gene Ontology (GO) terms [50] [23] associated with each of the protein interactors as binary features in a second type of dataset. There are three types of GO Terms, reflecting different types of information about a protein’s functions, namely: biological process, molecular function and cellular component. All 3 GO Term categories were used as predictive features in the dataset, for the sake of completeness.

Each binary feature indicates whether or not an instance (compound) is indirectly associated with a given GO term. More precisely, the feature value ‘1’ means that at least one of the proteins that interact with the compound (as recorded in the STITCH database) is annotated with the corresponding GO term. Conversely, the feature value ‘0’ means that none of the proteins interacting with the compound have been annotated with that GO term. The proteins’ GO term annotations were obtained using the *goatools* (version 1.1.6) Python library and data from the Gene Ontology version 1.4, downloaded in 11-2021 (Website: geneontology.org). The created dataset initially had 9000 unique GO terms, which were reduced to 7706 predictive features after applying the simple frequency-threshold filter.

#### 4.1.3 Phenotypes Dataset

In this dataset, each binary feature indicates whether or not a compound (instance) is indirectly associated with a given Phenotype Ontology (Physiology) term in the Wormbase database [24] (Website: wormbase.org/tools/ontology_browser). More precisely, the feature value ‘1’ means that at least one of the proteins that interacts with the compound (as recorded in STITCH) is annotated with the corresponding Phenotype Ontology term; otherwise, the feature takes the value ‘0’. Note that the Phenotype Ontology has physiology and anatomical phenotypes, and we only used the physiology terms, because anatomical characteristics are not as relevant for our prediction problem (whether or not a compound’s administration extends *C. elegan*’s lifespan). The created dataset initially had 1783 physiology phenotype features, which were reduced to 1272 after applying the frequency-threshold filter.

#### 4.1.4 Age-related Genes Dataset

In this dataset, each predictive feature indicates whether or not a compound (instance) interacts with a given age-related gene. To create these features, we used the lists of *C. Elegans* genes in the GenAge [51] [25] (Build 20, downloaded in 11-2021; Website: genomics.senescence.info/genes) and the GenDR [26] (Build 4, downloaded in 11-2021; Website: genomics.senescence.info/diet) databases.

The GenAge database is a collection of genes from different organisms known to be associated with longevity and/or ageing. The GenDR database is a collection of genes specifically associated with dietary restriction (including caloric restriction), included in the definition of this feature type because this intervention is commonly associated with lifespan increase in multiple organisms, and some of those genes were not listed in GenAge.

Based on the proteins coded by these genes, we defined the value of their feature using the list of protein interactors associated with each compound. If at least one of the interactors of a compound is coded by a given gene (feature), its feature value in the dataset is ‘1’, otherwise the value is ‘0’. The created dataset initially had 683 binary features in total, each referring to one gene in GenAge or GenDR. These were reduced to 365 features, after applying the frequency-threshold filter.

### 4.2 Filter Feature Selection Methods

Filter methods are a type of feature selection (FS) method used in a pre-processing phase of machine learning – before training the classification algorithm [52]. They calculate a score for each feature in the dataset, usually based on the distribution of its values in relation to the class label. Then, the top *k* (a user-specified parameter) features with the highest scores are kept, and all others are discarded. Note that filters are independent from the classification algorithm, in contrast to the more computationally expensive wrapper FS methods [52].

Filter methods differ mainly in terms of how they calculate the features’ scores, as there are various ways to measure feature importance, and no method is the best for all datasets. For our experiments in this study, we selected 5 different filter methods and, in addition to these, we developed a sixth (meta)-method called Auto-Filter, which automatically selects the best candidate filter method for each dataset using a data-driven approach. These filter methods are described next.

#### 4.2.1 Information Gain

This filter calculates the score of a feature as the value of the Information Gain (reduction of Entropy) obtained by partitioning the instances of a dataset into subsets, based on the values of that feature. Notably, this measure is known to be biased in favour of features with many values [53], but this is not an issue in our case as all predictive features in our datasets are binary. The Information Gain is calculated for each feature *F* as follows.

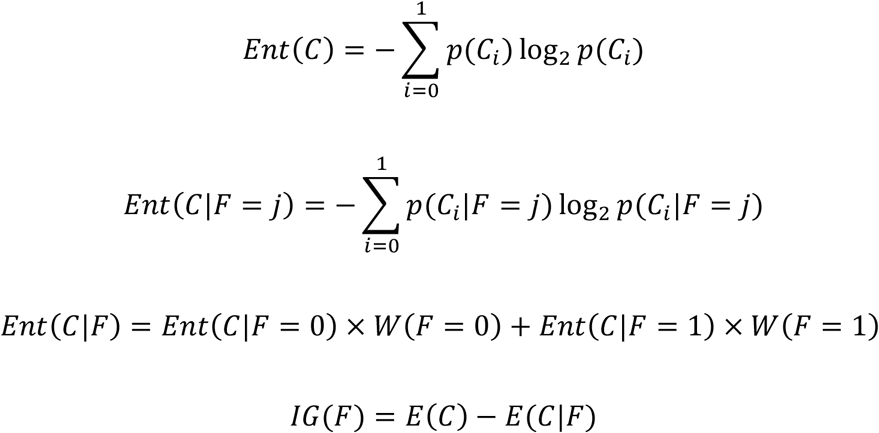

Where *Ent(C)* is the entropy of the class labels on the training data, *p*(*C*_*i*_) is the empirical probability (relative frequency) of class *i* in the training set (as our classification problem is binary, the class is either *0* or *1*), and *Ent(C*|*F)* is the entropy of the class labels conditioned on the values of feature *F* on the training data. We calculate the entropy of the class labels before and after splitting the dataset using the current feature, i.e. calculating *Ent(C)* and *Ent(C*|*F)* respectively, the latter being a weighted sum of the Entropies of the class labels in both splits (data subsets where *F = 0* and *F = 1*), where the weights are the proportions of instances with *F = 0* and *F = 1* in the training data. The information gain is the difference between the entropy *Ent(C)* and the conditional entropy *Ent(C*|*F)*, with larger values indicating a greater reduction of class-label entropy, i.e. a stronger predictive power associated with the feature.

#### 4.2.2 The Chi^2^ (Chi-Squared) Statistic

The Chi^2^ test is a statistical hypothesis test used to get an estimation of the degree of association between two categorical variables (in our case, each predictive feature and the class variable). It compares the observed and expected frequencies of each combination of values of those two variables, and larger differences between these values indicate that the variables have a stronger association [54]. We used the value of the Chi^2^ statistic calculated by this test as the score for a filter method, as the higher this value gets, the greater the association between the feature and the class variable. The Chi^2^ score is calculated as follows.

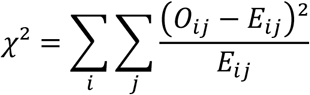

Where *O*_*ij*_ represents the observed frequency of the co-occurrence of the *i*-th value of a feature and the *j*-th class label, i.e. the number of instances with the *i*-th feature value and the *j*-th class label in the training set, and *E*_*ij*_ represents the expected value of that frequency of co-occurrence under the assumption that the feature and the class variable are statistically independent.

#### 4.2.3 Decision Stump Filters

A Decision Stump is the simplest version of a decision tree classifier, where the class label is decided based on a single node partition of the data, using a single feature. It usually does not have much predictive power on its own, but it can be used in other contexts such as providing a score for a filter method [55]. The score of a feature is calculated by training a decision stump classifier with a very narrow subset of the training data containing only that feature and the class variable, then evaluating the trained classifier on a subset of data which was not used for training (to estimate generalisation performance). In our experiments, the performance of a decision stump classifier was estimated by an internal 5-fold cross-validation procedure, applied to the training set only (i.e. not using the test set). Hence, the training set is divided into 5 folds and the decision stump classifier was trained 5 times, each time using a different fold as the ‘validation set’ (to estimate generalisation performance) and the other four folds as a ‘learning set’. We used the median of the five AUC values obtained by the decision stump classifier over the 5 validation sets as the score for this filter.

#### 4.2.4 Log Odds Ratio

The associations between two categorical variables, such as a binary predictive feature and the binary class variable in our datasets, may be displayed through a contingency table. In these tables each cell contains a count (*n*) or probability (*p*) of each combination of the values of the two variables, as shown in Figure 2. Some measures of association can be calculated based on this representation, including the Log Odds Ratio and the Asymmetric Optimal Prediction filters, described in this Subsection and the next.

**Figure 2.**
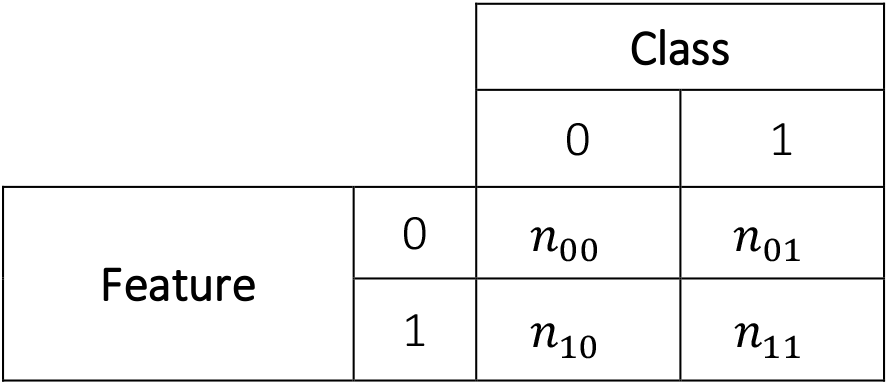
structure of a contingency table

The Odds Ratio [56] is a measure of association applicable to binary variables, which estimates the odds of an outcome based on the exposure (i.e., how much the odds of getting a ‘1’ value for a variable change based on the value of the associated variable). Odds values higher than 1 indicate an increased probability of success, and values lower than 1 indicate the opposite.

Smaller data samples cause the distribution of the Odds Ratio to be highly skewed. Thus, the natural logarithm of this measure, Log Odds Ratio, is used instead. The Log Odds Ratio between each feature and the class variable was used as a score for this filter method, calculated as follows.

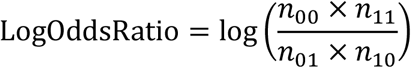

#### 4.2.5 Asymmetric Optimal Prediction

The Asymmetric Optimal Prediction (AOP) is another measure of association between two categorical variables, with the distinction of measuring an asymmetric predictive relationship, i.e. measuring to what extent the value of variable *A* can be well predicted by the value of another variable *B*, regardless of strength of the converse type of prediction (predicting *B* from *A*) [57]. This is relevant because in the classification task we use the feature value to predict the class label, not vice-versa, therefore it can be beneficial to focus on this asymmetrical association between the two variables.

The AOP measure compares two scenarios for predicting the class label for a randomly chosen instance: (1) knowing only the class-distribution in the training data and (2) knowing both the class distribution and the value of a feature *F*. The AOP measure is calculated using the probability of error in both cases, and increases in value as the probability of error in scenario 2 reduces. The AOP score is calculated as follows.

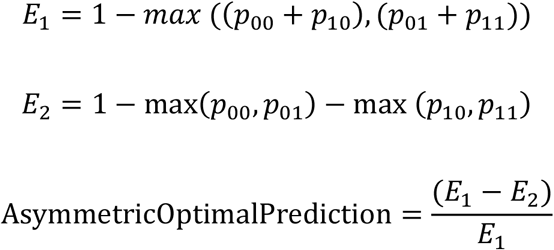

where *p*_00_ = *n*_00_/*n, p*_10_ = *n*_10_/n, *p*_01_ = *n*_01_/*n, p*_11_ = *n*_11_/*n*; *n*_00_, *n*_10_, *n*_01_ and *n*_11_ are as defined in Figure 2, *n* is the total number of training instances, *E*_*1*_ is the probability of a prediction error when an instance is predicted to have the most frequent class label among all training instances (i.e. the class prediction ignores the feature value), and *E*_*2*_ is the probability of a prediction error when an instance is predicted to have the most frequent class label among the training instances which have the same feature value (1 or 0) as the current instance (i.e. the class prediction is based on the feature value).

### 4.3 Filter Ensembles

As our datasets have imbalanced class distributions, the majority-class instances can skew the feature score values computed by a filter. One simple way to mitigate this would be to undersample the majority-class instances in the training set before calculating the filter scores, so that the data used by the filter is balanced [58]. However, this would cause most majority class instances to be completely ignored during the FS process.

Therefore, we applied instead a more robust way to calculate filter scores that also addresses the class imbalance issue, named Filter Ensembles. This strategy consists of using an ensemble of filters with bootstrap samples, combining their scores to get a final score value for each feature, which is calculated using only balanced datasets (i.e., each bootstrap sample has its majority class instances undersampled to a 1:1 ratio of positive- and negative-class instances). A similar strategy was used by [59] in the context of gene-gene interaction, and by [60] in the context of biomarker identification.

In our experiments we used an ensemble of 50 filters, but this number can be adjusted based on the available computational resources and the class distribution of the original dataset (more imbalanced datasets might need more filters). Each of the 50 balanced datasets is fed into a filter method to calculate the features’ scores, and the final score of a feature with that filter is the median value over these scores. This strategy is computationally expensive, but it can be implemented as a parallel algorithm, making use of multi-thread and multi-core architectures to reduce running time, as the scores for each of the 50 filters can be calculated simultaneously.

We performed a preliminary set of experiments comparing using the proposed Filter Ensembles strategy to not doing so (i.e., running a single filter using the full, unbalanced dataset), and concluded that the strategy led to classifiers with better predictive accuracy in general, and was therefore worthwhile. The discussion for this set of experiments is out of the scope of this paper, but the result tables detailing them are available in the Supplementary Material (Tables S1-S4).

### 4.4 Auto-K: Automatically selecting the number of top-ranked features to be kept

Generally, the user of a filter method needs to manually choose the number of top-ranked features to be selected by the method, denoted by *k*. Naturally, this choice can significantly impact the performance of a classifier trained with the selected features. In order to make our filter methods more adaptable, and reduce the impact of subjective user choices of *k* in the predictive performance of classifiers, we used an automated process for selecting the best *k* value for a dataset out of a set of candidate *k* values, which we named Auto-K.

The candidate *k* values were defined based on the numbers of features in the original datasets, as follows. For the Interactors and GO terms datasets, which have 5788 and 7706 features respectively, we set the candidate *k* values as 250, 500, 750, 1000. For the Phenotypes dataset, which has 1272 features, we set the candidate *k* values as 100, 200, 300, 400. Finally, for the GenAge/GenDR dataset, which has 365 features, we set the candidate *k* values as 50, 100, 150, 200.

Auto-K works as follows. For each fold of the 10-fold external cross-validation, an internal 5-fold cross-validation is used to train RF classifiers using each candidate *k* value, and the value that results in models with the highest median AUC is chosen. Note that only the training data is used by the Auto-K method, as the test data cannot be accessed prior to evaluating the final classifier.

### 4.5 The Auto-Filter Method for Feature Selection

As the performance of a filter method depends largely on the data distribution, making an automated data-driven choice of the best filter method for each dataset intuitively should lead to better predictive performance, compared to making a fixed choice regardless of the data.

Thus, we implemented another filter approach named the Auto-filter approach. In addition to automatically selecting the best *k* value for the number of top-ranked features to be kept in the dataset, the Auto-Filter approach also selects the best candidate filter method out of the 5 candidate filters discussed in Section 4.2 or out of their filter ensemble counterparts (Section 4.3). This automated filter-method selection is based on an internal 5-fold cross-validation process applied to the training set, where it trains RF classifiers using each of the possible 15 combinations of a filter (or filter ensemble) method and a *k* value (5 filters times 3 *k* values).

The median AUC (Area Under the ROC curve) of the classifiers over the 5 folds of the internal cross-validation is used to select the best filter or filter ensemble method (the one with the highest median AUC), with the AUC variance as a tie-breaking criterion (the lower the variance, the better). As the Auto-Filter procedure is used inside an external cross-validation process, it will be run once for each fold in that external process, using each training dataset. Note that different candidate filter or filter ensembles and *k* values might be selected across the different folds of the external cross-validation. The pseudocode in Algorithm 1 represents the Auto-Filter procedure used in each fold of the external cross-validation.

#### Algorithm 1

The Auto-Filter procedure. It receives a set of candidate filter methods *S*, a set of candidate *k* values *K*, the training dataset of the external cross-validation, a classification algorithm (in this work, random forest) to be used in the internal comparisons of the candidate filters, and a target predictive performance metric (in this work, the AUC). It returns the candidate filter and *k* value combination with the largest average score. Note that in this pseudocode the term ‘filter method’ is being used in a generic way, it can denote either a single filter method or its counterpart filter ensemble.

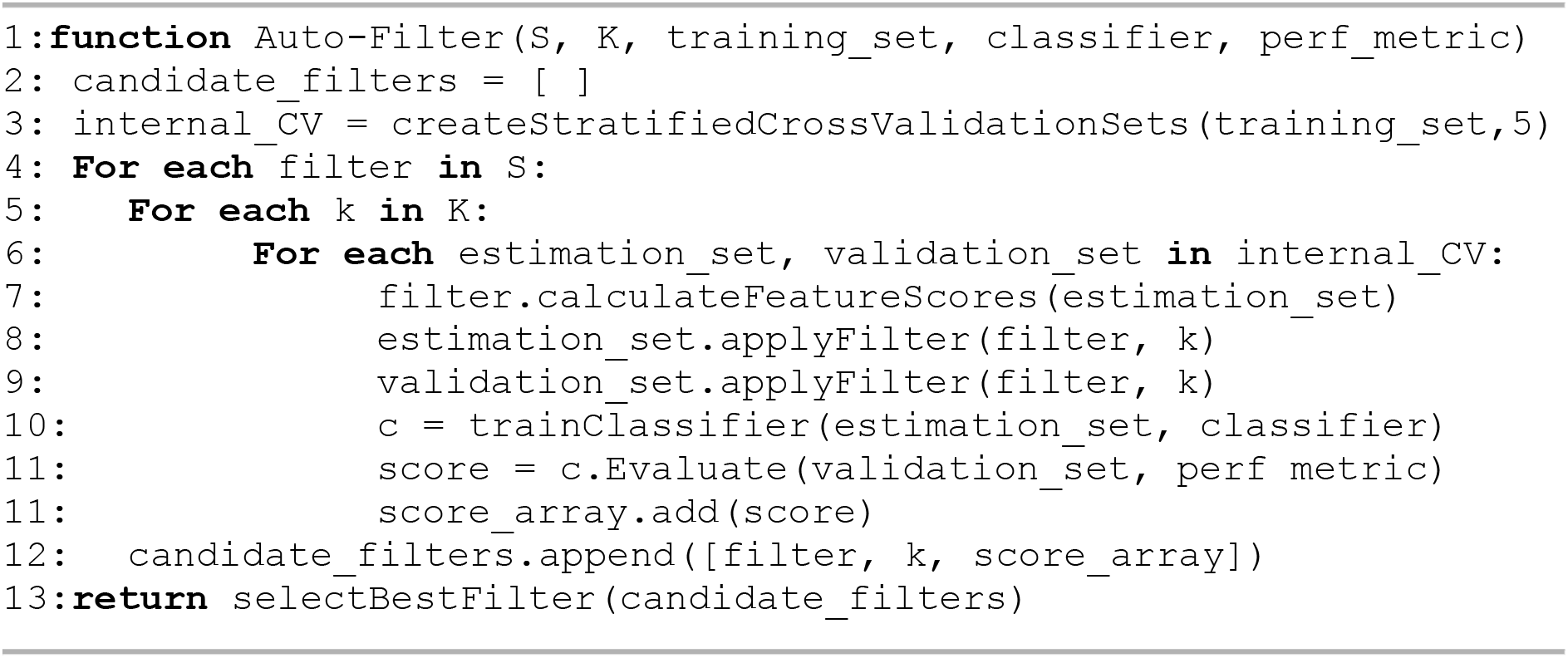

The Auto-Filter approach is flexible, as the user may select any number of candidate filters (with or without using filter ensembles) and *k* values, as well as the classification algorithm and performance metric used in its internal method for selecting the best candidate filter method (e.g., it might be more relevant to optimise the filter choice using the F-Score or the AUC metric, depending on the characteristics of the project). However, the Auto-Filter approach has the disadvantage of being computationally costly, as it requires many runs of each candidate filter and the classification algorithm. This disadvantage can be alleviated by a parallel implementation, as most parts of the process (i.e., calculating the scores of each candidate filter, the folds in the internal cross-validation, and the external cross-validation process) are independent.

### 4.6 Experimental Setup

In order to test the feature selection approaches described in this paper, we ran experiments comparing Random Forest (RF) classifiers trained using each of them. In all experiments we performed a 10-fold cross-validation process and report the median value of the well-known Area Under the Receiver Operating Characteristic curve (AUC) [61]. The RFs were trained with 500 trees and the number of features randomly sampled as candidate features for each node was 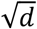 (rounded up to nearest integer, with 0.5 being rounded up), with *d* being the number of features in the current dataset.

As the datasets created for this research have a class-imbalance issue (22.6% of positive-class instances and 77.4% of negative-class instances), we trained our RFs using the Balanced Random Forest (BRF) method [31]. The BRF method draws a bootstrap sample of minority class instances for each tree in the forest, and randomly draws the same number of instances from the majority class instances, meaning the subset of instances used to generate each decision tree has a balanced ratio (1:1) of instances from each class.

## 5 Acknowledgements

We thank Dr. Taravat Ghafourian (University of Bedfordshire, UK) for her help in dataset preparation, in particular for checking the matching between some compound names in the DrugAge and STITCH databases. This project was funded by a research grant from the UK’s Biotechnology and Biological Sciences Research Council (BBSRC), grant reference numbers BB/V007971/1 and BB/V010123/1, to AAF and JPM. DrugAge is supported by a Biotechnology and Biological Sciences Research Council UK (BB/R014949/1) grant to JPM.

## 6 Author Contributions

JPM and AAF conceived the overall project. CR and AFF designed the machine learning methodology used in the experiments. All authors designed the structure of the created datasets. CR created the datasets, implemented all required machine learning algorithms and ran all the computational experiments. CR and AAF analysed the predictive accuracy results. CF and JPM analysed the biological results. The manuscript was written mainly by CR, but all authors contributed to writing and revising the manuscript.

## 8 Data Availability

The datasets used in the experiments and the program code for the feature selection methods will be made freely available on the web when the paper is published.

