## Supplementary Tables S1-S4 for "Predicting Lifespan-Extending Chemical Compounds with Machine Learning and Biologically Interpretable Features"

### Supporting Information

#### Supplementary Tables – Selecting the best strategy for feature selection

Supplementary Tables S1 through S4 compare the median AUC values obtained by the Random Forest algorithm when using two different versions of each candidate filter method, with the highest value highlighted in boldface. The first version, named single filter, simply applies the standard filter method without using any ensemble or class balancing approach. Hence, it computes scores for the features using the dataset's original imbalanced form, where the majority class will usually have a larger impact on the score of a feature. The second version, named filter ensemble, addresses that issue by computing scores using an ensemble of balanced filters as proposed in Section 4.3 of the main paper. In order to determine whether the significant computational cost added by using the second type of method is worthwhile, we compared these two types of filter methods in a set of experiments using the 4 datasets prepared in this work, each dataset using a different type of predictive feature, as specified in the titles of Tables S1 through S4. The last row of each table shows the AUC value for the baseline approach of simply training the classifier using the full set of features, without performing any feature selection in a pre-processing phrase.  Note that, in all experiments, the value of *k* (the number of features selected by a filter or filter ensemble method) is automatically selected by the Auto-K method described in Section 4.4 of the main paper.

*Table S1. Median AUC values obtained by Random Forest using the single filter vs filter ensemble methods in a pre-processing phase – Protein Interactors dataset.*

| **Interactors Dataset** | **Single Filter** | **Filter Ensemble** |
|---|---|---|
| Information Gain | 0.736 | 0.743 |
| $Chi^2$ | 0.730 | 0.738 |
| Decision Stump (GMean) | 0.731 | 0.738 |
| Asymmetric Optimal Prediction | 0.724 | 0.742 |
| Log Odds Ratio | 0.725 | 0.713 |
| Auto-Filter | 0.759 | **0.801** |
| Baseline (no filter method) | 0.717 | |

*Table S2. Median AUC values obtained by Random Forest using the single filter vs filter ensemble methods in a pre-processing phase – GO Terms dataset.*

| **GO Terms dataset** | **Single Filter** | **Filter Ensemble** |
|---|---|---|
| Information Gain | 0.753 | 0.744 |
| Chi² | 0.740 | 0.725 |
| Decision Stump (GMean) | 0.719 | 0.734 |
| Asymmetric Optimal Prediction | 0.725 | 0.750 |
| Log Odds Ratio | 0.721 | 0.708 |
| Auto-Filter | 0.779 | **0.818** |
| Baseline (no filter method) | 0.767 | |

*Table S3. Median AUC values obtained by Random Forest using the single filter vs filter ensemble methods in a pre-processing phase – Physiology Phenotypes dataset.*

| **Phenotypes Dataset** | **Single Filter** | **Filter Ensemble** |
|---|---|---|
| Information Gain | 0.759 | 0.722 |
| Chi² | 0.711 | 0.729 |
| Decision Stump (GMean) | 0.714 | **0.761** |
| Asymmetric Optimal Prediction | 0.755 | 0.741 |
| Log Odds Ratio | 0.724 | 0.706 |
| Auto-Filter | 0.719 | 0.728 |
| Baseline (no filter method) | 0.741 | |

*Table S4. Median AUC values obtained by Random Forest using the single filter vs filter ensemble methods in a pre-processing phase – GenAge/GenDR dataset.*

| **GenAge/GenDR Dataset** | **Single Filter** | **Filter Ensemble** |
|---|---|---|
| Information Gain | 0.709 | 0.727 |
| Chi² | 0.740 | 0.751 |
| Decision Stump (GMean) | **0.757** | 0.742 |
| Asymmetric Optimal Prediction | 0.739 | 0.744 |
| Log Odds Ratio | 0.708 | 0.721 |
| Auto-Filter | 0.702 | 0.725 |
| Baseline (no filter method) | 0.683 | |

As can be observed in these tables, in 3 out of the 4 datasets, the best AUC value (highlighted in boldface in each table) was obtained by the filter ensemble approach. In addition, in total, over the

24 pairs of results comparing single filter vs filter ensemble (6 comparisons per table times 4 tables) models, the latter won in 16 (67%) of the cases. Hence, overall the filter ensemble approach performed better than the single filter approach.

After deciding to apply the filter ensemble strategy, we then compared the filter ensembles' results to determine the best FS method (regarding predictive accuracy) out of our set of 6 candidate filter ensemble methods. The proposed Auto-Filter method (described in Section 4.5 of the main paper) got the best median AUC results for two datasets, namely the Protein Interactors and the GO Terms datasets, with these two models being the two best models (i.e. with the two highest AUC values) overall. For the Physiology Phenotypes dataset, the best method was the Decision Stump filter ensemble, and for the GenAge/GenDR dataset the winner (among filter ensemble methods) was the $Chi^2$ filter ensemble. As the proposed Auto-Filter approach has the advantages of flexibility and robustness (since it selects the best filter ensemble for each dataset), and got the best results for two of the four datasets, we have chosen it as our main approach.
